# Sight-line hypothesis explains facial color patterns in terns and allies

**DOI:** 10.64898/2026.03.25.714058

**Authors:** Masaru Hasegawa

## Abstract

Conspicuous coloration in animals is generally thought to evolve and be maintained through inter- or intraspecific interactions such as mate choice, but this might not always be the case. The sight-line hypothesis proposes that conspicuous light-dark contrast in front of the eyes (hereafter, eyeline) evolves and is maintained due to viability selection, enhancing an individual visual acuity and thus evolutionarily associated with a particular foraging behavior that requires accurate aiming. However, empirical evidence that supports the sight-line hypothesis is virtually absent, with no studies demonstrating the key prediction that the direction of eyelines matters. Here, I tested the sight-line hypothesis using macroevolutionary analyses in terns and allies, which are a suitable study system, because they have variation in facial color patterns, including presence/absence and, if any, various angles of eyelines. They also have a large variation in foraging behavior, including picking, plunge diving, and skimming. As predicted by the sight-line hypothesis, tern lineages that require accurate aiming at foraging (e.g., plunge diving) are more likely to have eyelines. In addition, the evolutionary transition to the state with eyelines and these foraging behaviors was more likely to occur than the reverse transition. Furthermore, as expected by the fact that the direction of travel is upwardly deviated from the direction of the bills during skimming, the eyeline angle from bills was evolutionarily positively associated with the occurrence of skimming behavior. To my knowledge, the current study is the first to demonstrate that the direction of the eyeline matters, thereby strongly supporting the sight-line hypothesis.

---

Conspicuous coloration in animals is generally considered to have evolved and been maintained through interspecific interactions (e.g., as aposematic signals or concealment against colorful backgrounds) or intraspecific interactions (e.g., as sexual signals; reviewed in Andersson 1994). Bird plumage coloration would be a representative study system, demonstrating the importance of interactions with others (e.g., see Hill & McGraw 2006 for a review). However, inter- and intraspecific interactions would not be responsible for the evolution and maintenance of conspicuous coloration in some cases. Conspicuous facial coloration can be used to improve their own vision; for example, the black facial mask of predatory animals can have a function to reduce glare and thus improve foraging success (e.g., Bortolotti 2006; Yosef et al. 2012; Vrettos et al. 2021).

The sight-line hypothesis provides another explanation for conspicuous facial coloration (Ficken & Weillmot 1968; Ficken et al. 1971). This hypothesis proposes that animal eyelines can be utilized as an aiming sight (i.e., sight-line) to catch prey. This hypothesis has mistakenly been regarded as a variation of the black facial mask to reduce glare (e.g., Bortolotti 2006; see above) and thus has almost been forgotten, but, in really, this hypothesis stresses the importance of dark and light contrast in front of the eyes that assists fast and accurate aiming. In fact, the recent macroevolutionary analysis of hirundines, i.e., hyper-aerial insectivores, Hasegawa (2023) revealed that hirundines foraging on large (and thus active) flyers, which thus require accurate aiming, are more likely to have eyelines than those foraging on small (and thus planktonic) aerial insects, thereby supporting the hypothesis. Moreover, Hasegawa (2023) demonstrated that having dark facial coloration alone does not explain the observed pattern, thereby emphasizing the importance of dark–light contrast in front of the eyes. However, except for the macroevolutionary analysis of hirundines (Hasegawa 2023), empirical evidence that supports the sight-line hypothesis is virtually lacking, at least in the modern scientific framework. In particular, under the sight-line hypothesis, the direction of the eyeline should align with the line of sight and thus be associated with foraging behavior (Ficken et al. 1971), which remains untested.

Here, I tested the sight-line hypothesis using terns, i.e., “sea-swallows” (Gochfeld & Burger 1996), including skimmers and noddies (thus, terns and allies, which I call terns hereafter for brevity; see Winkler et al. 2020; Cerný & Natale 2022 for their phylogenetic relationships within Charadriiformes). Similar to hirundines, terns are a monophyletic clade highly adapted to aerial locomotion with long wings and short legs (Gochfeld & Burger 1996). Terns provide a suitable opportunity to evaluate the sight-line hypothesis, because of interspecific variation in facial color patterns (e.g., presence/absence of eyelines: Bridge et al. 2005) and in foraging behavior (Gochfeld & Burger 1996). They adopted visually-guided foraging (e.g., Martin et al. 2007; Luca et al. 2024). Some terns adopt foraging behaviors that need fast and accurate aiming, including plunge diving for fish and other aquatic prey (Gochfeld & Burger 1996; Fig. 1). Other terns adopt foraging behaviors that does not require fast and accurate aiming, for example, seizing insects from surface of ground (Gochfeld & Burger 1996), thereby providing an opportunity to test the evolutionary association between accurate aiming at foraging and the presence/absence of eyelines. Furthermore, another foraging behavior—namely, skimming, in which birds fly low over the water with the lower bills partially submerged to swiftly catch fish and other prey—can be found in this clade (e.g., the black skimmer, *Rynchos niger*: Winkler et al. 2020). During skimming, terns need to look forward from their bill tips toward the direction of travel (i.e., positive angles from the bills), because bills, when overlapped with prey, interfere with the direct sighting of prey, and because birds require accurate aiming of prey before manipulating bills to seize prey (Fig. 1; avoiding obstacles before collision is also important; Martin et al. 2007). Thus, the eyeline angle (Fig. 2) would be evolutionarily positively associated with the occurrence of skimming behavior under the sight-line hypothesis.

**Fig. 1.**
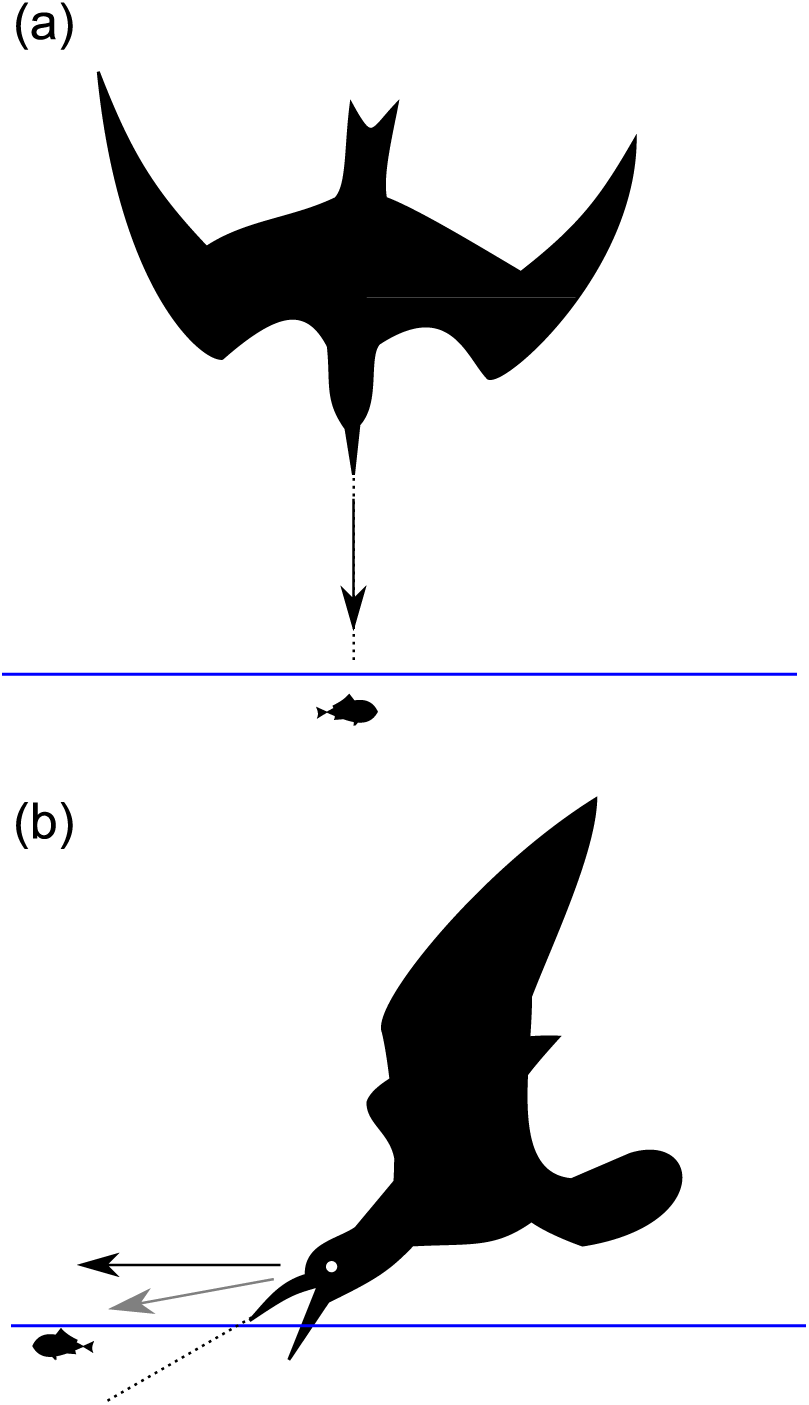
Two different foraging behaviors: a) plunge diving; b) skimming. Note that the direction of travel (shown in a black arrow) is positively deviated from the direction of bills (shown in a dotted line) in skimming, which would not always be the case in plunge diving. Blue lines indicate the water surface. During skimming, terns should pinpoint potential prey items before their bills interfere with direct sighting of prey (i.e., the line of sight, shown in gray arrow, should be anywhere between direction of travel and direction of upper bill), which contrasts with plunge diving, in which the sight-line can deviate in either a positive or negative direction when they aim at prey before diving. In panel (b), bills are opened, which illustrates the actual skimming posture, and thus the line of sight would be further deviated upwardly from the direction of closed bills, from which the eyeline angle was measured (see text).

**Fig. 2.**
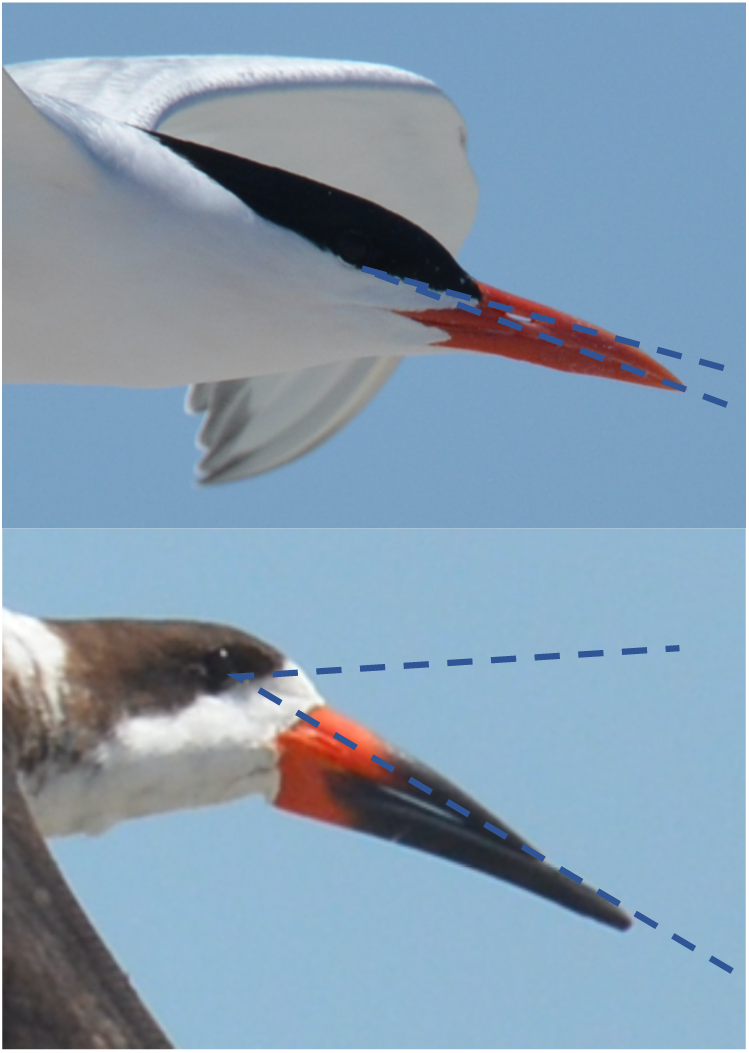
Examples of eyelines: upper panel) the royal tern, *Sterna maxima*; lower panel) the black skimmer, *Rynchos niger*. Note the difference in eyeline angles, defined as the angle from eye-to-bill tips to eyelines (see Methods), shown in lines.

Using phylogenetic comparative analyses of terns, I firstly tested whether or not species with foraging that requires accurate aiming are more likely to have eyelines than others. I also conducted an analysis of evolutionary pathways to clarify the transition patterns of the presence/absence of eyelines in relation to foraging that requires accurate aiming or not. Furthermore, I here tested the key prediction of the sight-line hypothesis that the direction of eyelines is evolutionarily associated with foraging behaviors through differential lines of sights. I predicted here that the occurrence of skimming behavior would be evolutionarily positively associated with the eyeline angle (see above). I found patterns consistent with these predictions, thereby supporting the sight-line hypothesis.

## Methods

### Data collection

Information on the presence/absence of eyelines (i.e., contrasting lines projected from eyes to forehead) and bill coloration was obtained from illustrations given in “The Birds of the World” (Winkler et al. 2020). Any dark–light color contrasts in front of the eye were regarded as eyeline (e.g., color, length, and width; Ficken & Weillmot 1968; Ficken et al. 1971), but most of them were long black–white borderlines (e.g., black “cap” on white below). When separate illustrations were available for breeding/nonbreeding plumage, the breeding illustration was chosen, partially because nonbreeding appearance could not be obtained for some species (e.g., the black-naped tern, *Sterna sumatrana*) and because breeding color pattern was clear-cut and thus was more easily measurable than the nonbreeding one. When eyelines were present in the focal species illustration, the angle from eye-to-bill tips to eyelines were measured using ImageJ software. The angle was measured twice per species, and the mean values were used for analyses. Repeatability of the two measurements was high (repeatability = 0.98, F_39,40_ = 134.89, *P* < 0.0001; Lessels & Boag 1987).

In contrast to hirundines, in which all species have black bills (Hasegawa 2023), some tern species have colorful bills (e.g., red, orange, yellow) instead of black bills, which can affect facial color pattern (e.g., see Minias & Janiszewski 2020 for gulls, a sister clade of terns). For example, colorful bills might have evolved via sexual (and social) selection, and hence those species with colorful bills might experience different selection on facial color patterns compared with those with black bills. Thus, I took account of bill coloration as a dichotomous variable (colorful or black) when analyzing evolutionary associations between eyelines and foraging behaviors. This potential influence was controlled out by including them as a covariate (see statistics section for details).

Information on foraging behavior was obtained from the same literature (Winkler et al. 2020) to distinguish those that mainly use foraging that requires accurate aiming or not. The former group includes those using “diving” (or “plunge diving”) and “skimming” (i.e., those that require swift and accurate aim), whereas the latter group doesn’t include these categories (and instead uses picking, dipping, and other foraging behaviors; note that we included catching insects in this category). Information on body size (as a continuous measurement) and migratory habit (dichotomous category, migratory or not, based on whether or not they had separate breeding sites from wintering sites, which is shown in yellow in the distribution maps) was obtained from Gochfeld & Burger (1996), except for the little white tern *Gygis microrhyncha*, which is not listed in the literature and thus Winkler et al. (2020) was used instead for information on this species.

The occurrence of skimming in each species, which is relatively minor behavior except for skimmers (note that skimmers are the only species that mainly use skimming), was searched for using the same literature listed above (Gochfeld & Burger 1996; Winkler et al. 2020), along with additional literature (Supplementary Material 1 for detailed information). To focus on skimming at foraging (i.e., in which sight-line should be important; see Introduction), skimming for other uses was discarded (e.g., when drinking water or washing bills, which do not require aiming at prey items). In addition, information on European and North American terns are abundant compared with those in other regions (e.g., Snow & Perrins 1998; Larsson & Olsen 2010; note that “the Birds of the World” mainly consists of “Handbook of the birds of the world” and “Birds of the North America,” as is evident from revision history of “The Birds of the World”; Winkler et al. 2020). Thus, to account for the differential information availability, tern species were grouped as European and North American, and others using Larsson & Olsen (2010) and additional information on skimmers, which are not listed in Larsson & Olsen (2010), is obtained from Winkler et al. (2020). Finally, of 48 listed tern species in “The Birds of the World,” the description of one species, the Chinese crested tern, *Sterna bernsteini*, is unavailable (Winkler et al. 2020), and thus the final sample size was 47. Table S1 shows the dataset for the current study.

### Statistics

A Bayesian phylogenetic mixed model with a binomial error distribution was used to investigate the presence/absence of eyelines. In addition to foraging behavior (i.e., foraging that requires accurate aiming or not; see above), I included some confounding variables, which might be associated with plumage and ecological traits, as is the case in hirundines (i.e., body size and migratory habit; Hasegawa 2023). Bill coloration was included as an additional confounding factor (see previous section). To account for phylogenetic uncertainty, models were fit to each tree and applied multimodel inference using 1000 alternative trees for terns from birdtree.org (Garamszegi and Mundry 2014). Mean coefficients, 95% credible intervals (CIs) based on the highest posterior density, and Markov chain Monte Carlo (MCMC)-based *P*-values (P_MCMC_) were derived, together with the phylogenetic signal (de Villemereuil and Nakagawa 2014; also see Nakagawa and Schielzeth 2010 for detailed calculation). All analyses were conducted in R version 4.4.2 (R Core Team 2024) using the function “MCMCglmm” in the package “MCMCglmm” (version 2.36: Hadfield 2010). As before (e.g., Hasegawa 2023), a Gelman prior was used for the fixed effects while standardizing each continuous variable using “gelman.prior” in the package “MCMCglmm.” I ran the analysis for 140,000 iterations, with a burn-in period of 60,000 and a thinning interval of 80 for each tree. Similarly, to investigate the occurrence of skimming behavior in relation to the eyeline angle, another Bayesian phylogenetic mixed model with a binomial error distribution was built. Because I focus on the initial evolution of skimming behavior rather than its fixation as a major foraging behavior (which is impractical to analyze because only one clade, skimmers, has skimming behavior as a major foraging behavior; see above), we predicted that terns with more positive angles of eyelines were more likely to have skimming behaviors than others. In other words, adopting skimming behaviors as only a minor foraging behavior would not be a major selection force to change the eyeline angles, and thus the eyeline angle as a dependent variable would be inappropriate. The eyeline angle was log-transformed, i.e., log (eyeline angle + 10), to better fit the model (Shapiro–Wilk normality test: before: W = 0.80, *P* < 0.0001; after: W = 0.96, *P* = 0.22). This time, in addition to three potential confounding factors noted above (i.e., body size, migratory habitat, and bill coloration), I also included distribution range (i.e., North American & Europe or not) as another potential confounding factor, because the research progress might affect the description of relatively minor behavior (skimming, here).

In addition, as in the previous study conducted in hirundines (Hasegawa 2023), the discrete module in BayesTraits (Pagel 1994, 1999) was used to investigate evolutionary transitions among states with the presence/absence of eyelines and foraging behavior (i.e., foraging that requires accurate aiming or not). I used the same protocol as the previous study of hirundines (Hasegawa 2023). In brief, 1000 alternative trees of terns from birdtree.org were used, which was sufficient to control for phylogenetic uncertainty (Rubolini et al. 2015). Using MCMC methods in BayesTraits, the model was run for 1,010,000 iterations, in which a phylogenetic tree was chosen randomly from 1000 trees in each iteration, with a burn-in period of 10,000 and a thinning interval of 1000.

The Bayes factor (BF) was calculated by comparing the marginal likelihood of a dependent model that assumed correlated evolution of the focal two traits to that of an independent model assuming that the two traits independently evolved using the stepping stone sampler implemented in BayesTraits, as indicated by BayesTraits Manual V.3. Bayes factors of >2, 5–10, and >10 indicate positive, strong, and very strong evidence of correlated evolution, respectively. Note that the range of transition rate is 10e-32 to 100 (not 0 to 1), as explained in BayesTraits Manual V.3. I denoted means as the representatives of each transition rate. Likewise, I showed mean differences in transition rates, and P_MCMC_ values as the posterior probability that the differences would be higher (or lower) than 0. The reproducibility of the MCMC simulation was confirmed by calculating the Brooks–Gelman–Rubin statistic (Rhat), which should be <1.2 for all parameters (Kass et al. 1998) after repeating the analyses three times. To visually check replicated co-distribution (Maddison and FitzJohn 2016), examples of ancestral character reconstruction were presented using the functions “ace” in the R package “ape” and “plotTree” in the R package “phytools” (Revell 2012). I confirmed that statistical results supporting correlated evolution (see Results) did in fact represent replicated co-distributions throughout hirundines and thus false positives due to a few influential evolutionary events would be unlikely, as explained in other comparative studies (e.g., Clark et al. 2018).

## Results

### Eyeline and foraging that requires accurate aiming

Eyelines are widely distributed within the clade of terns (40 out of 47 species, 85%) and have been lost multiple times (Fig. 3). The interspecific distribution of eyelines closely matched the distribution of the evolutionary loss of foraging behaviors that requires accurate aiming (Fig. 3). In fact, when controlling for phylogeny and covariates (i.e., body size, migratory habit, and bill coloration), the probability of having eyelines was significantly explained by foraging behavior (Table 1): terns with foraging that requires accurate aiming were more likely to have eyelines. In addition, larger terns were found to be more likely to have eyelines (Table 1).

**Fig. 3.**
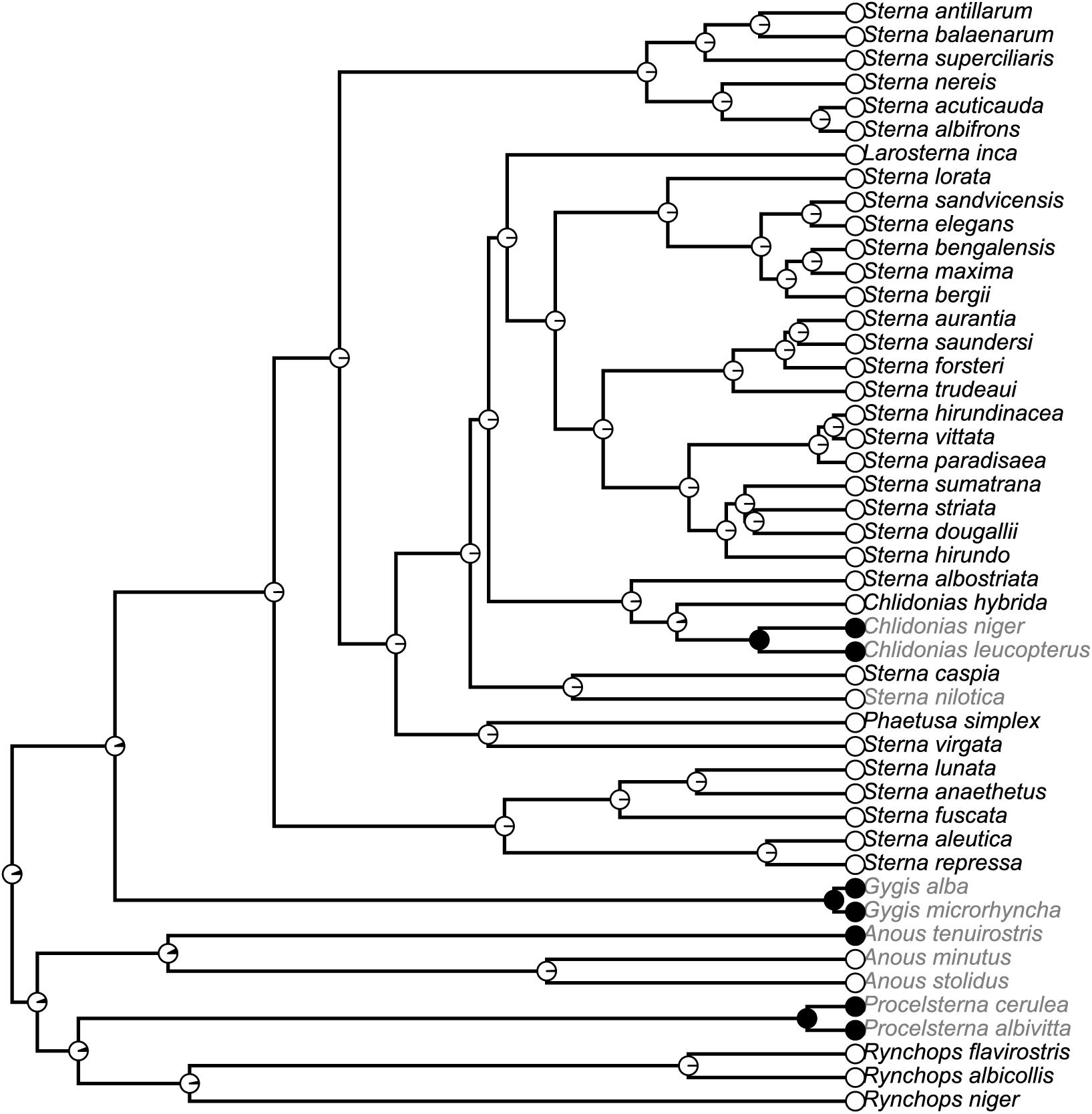
An example of ancestral character reconstruction of the presence/absence of eyelines in terns and allies using the function “plotTree” in the R package “phytools” (Revell 2012). White and black circles in tips indicate birds with and without eyelines, respectively. Similarly, the proportions of white and black in nodes represent the probability of an ancestral state with and without eyelines, respectively. Species names in black and gray correspond to species with and without foraging that requires accurate aiming.

**Table 1.**
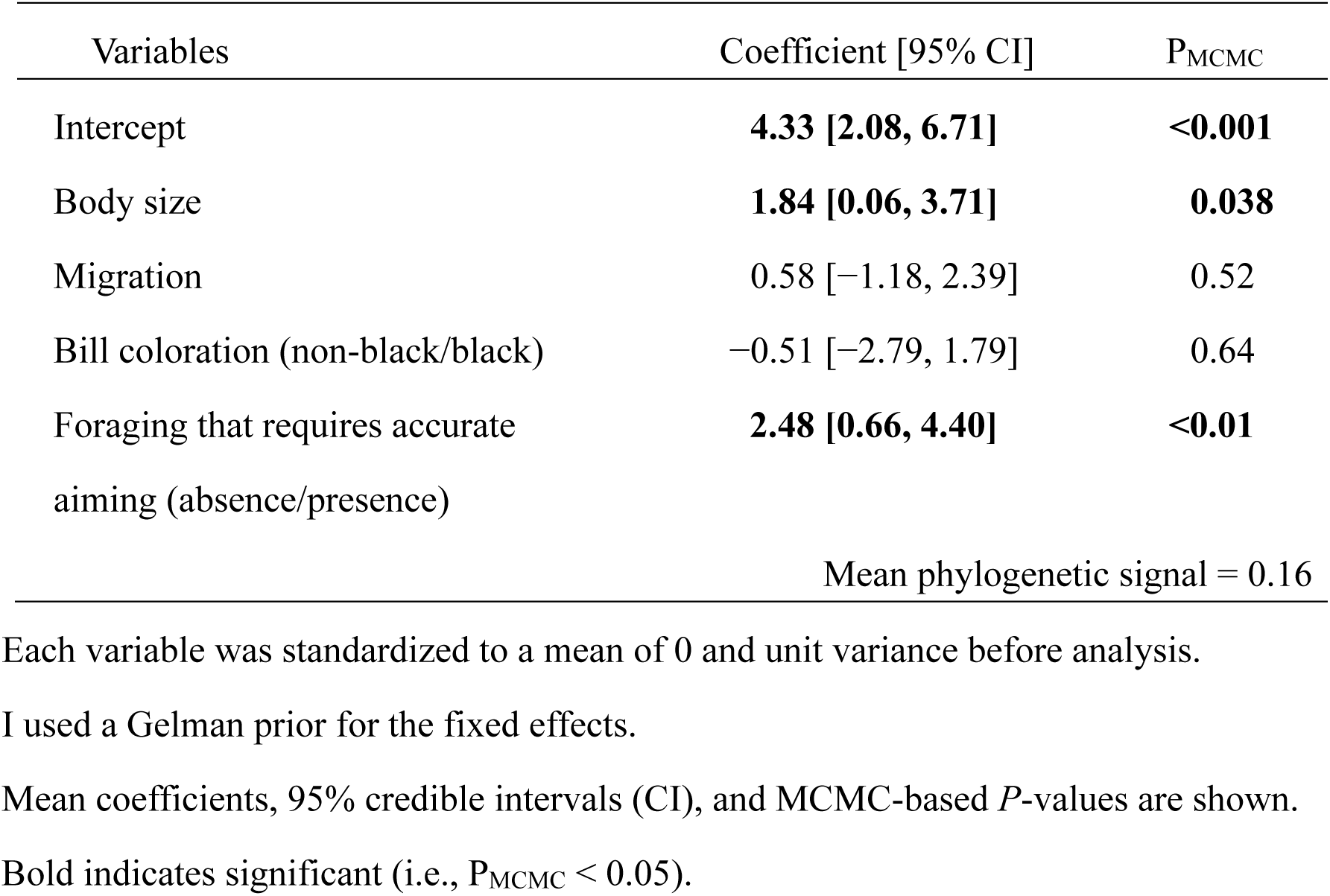
Multivariable Bayesian phylogenetic mixed model with a binomial error distribution predicting the presence/absence of eye-line in relation to foraging behavior in terns and allies (N = 47)

| Variables | Coefficient [95% CI] | P<sub>MCMC</sub> |
| --- | --- | --- |
| Intercept | **4.33 [2.08, 6.71]** | **<0.001** |
| Body size | **1.84 [0.06, 3.71]** | **0.038** |
| Migration | 0.58 [-1.18, 2.39] | 0.52 |
| Bill coloration (non-black/black) | -0.51 [-2.79, 1.79] | 0.64 |
| Foraging that requires accurate aiming (absence/presence) | **2.48 [0.66, 4.40]** | **<0.01** |
| Mean phylogenetic signal = 0.16 |  |  |

Each variable was standardized to a mean of 0 and unit variance before analysis. I used a Gelman prior for the fixed effects. Mean coefficients, 95% credible intervals (CI), and MCMC-based *P*-values are shown. Bold indicates significant (i.e., P<sub>MCMC</sub> < 0.05).

Analysis of evolutionary pathway using two state variables (i.e., presence/absence of eyelines and of foraging that requires accurate aiming) showed very strong support for correlated evolution of eyeline and foraging that requires accurate aiming (BF = 19.23). Transition to the state with eyelines and foraging that requires accurate aiming was more likely to occur than the reverse transition (mean difference = 54.09, 95% CI = 22.74, 81.72, P_MCMC_ < 0.01; Fig. 4).

**Fig. 4.**
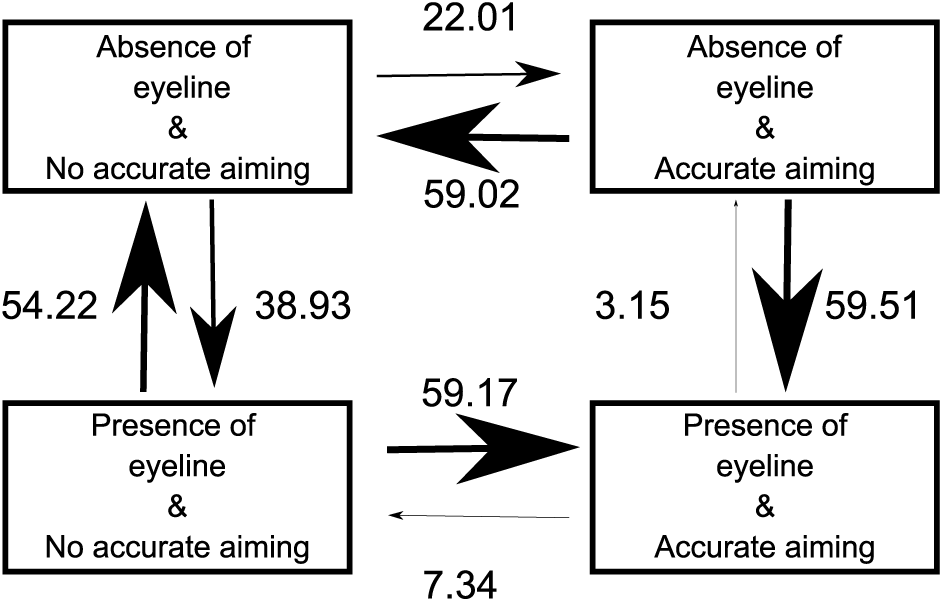
The most likely evolutionary transition between the presence and absence of eyelines in relation to foraging type (foraging that requires accurate aiming or not; see text) in the subfamily Sterninae. Model-averaged transition rates, which are reflected by arrow size, are depicted.

#### Eyeline angle and skimming

I tested whether or not the eyeline angle was associated with the evolution of skimming behavior. Skimming has evolved/lost multiple times, and 15 out of 40 (38%) species with eyelines demonstrated notable skimming behaviors (Fig. 5). The probability of having notable skimming behavior was significantly explained by the eyeline angle (Table 2) when controlling for phylogeny and covariates (i.e., body size, migratory habit, bill coloration, and region). Terns having eyelines with larger angles from bills were more likely to exhibit skimming behavior (Fig. 6). No other variables were significant (Table 2).

**Fig. 5.**
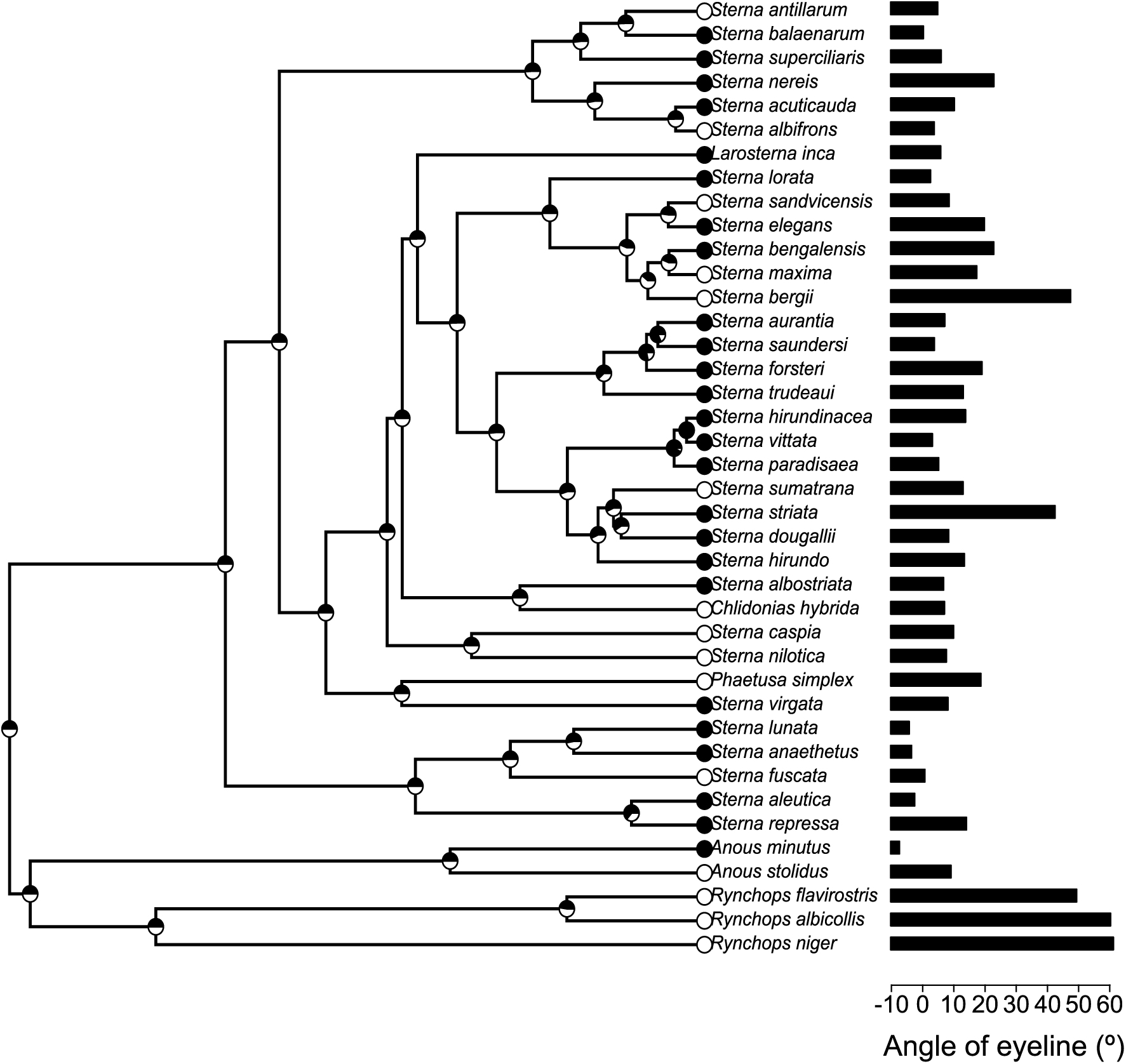
An example of ancestral character reconstruction of the occurrence of skimming foraging in terns (Aves: Sterninae) with eyelines using the function “plotTree” in the R package “phytools” (Revell 2012). White and black circles in tips indicate birds with and without the observation of skimming, respectively. Similarly, the proportions of white and black in nodes represent the probability of an ancestral state with and without skimming, respectively. Besides species names, the eyeline angle for each species is presented.

**Fig. 6.**
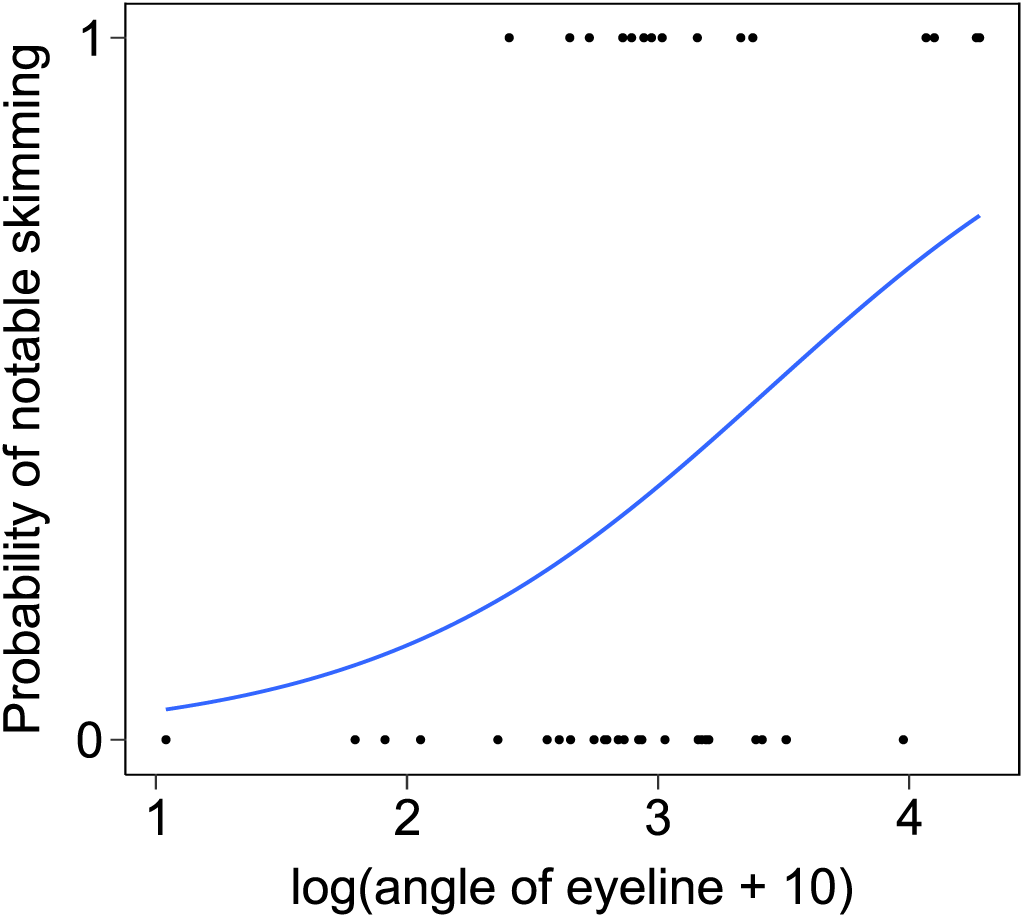
Terns with higher eyeline angles were predicted to have detectable skimming behaviors at a higher probability than others. Black dots indicate the presence/absence of detectable skimming behavior, and the simple logistic regression line that predicts the probability of notable skimming behavior. Formal test statistics are given in Table 2.

**Table 2.**
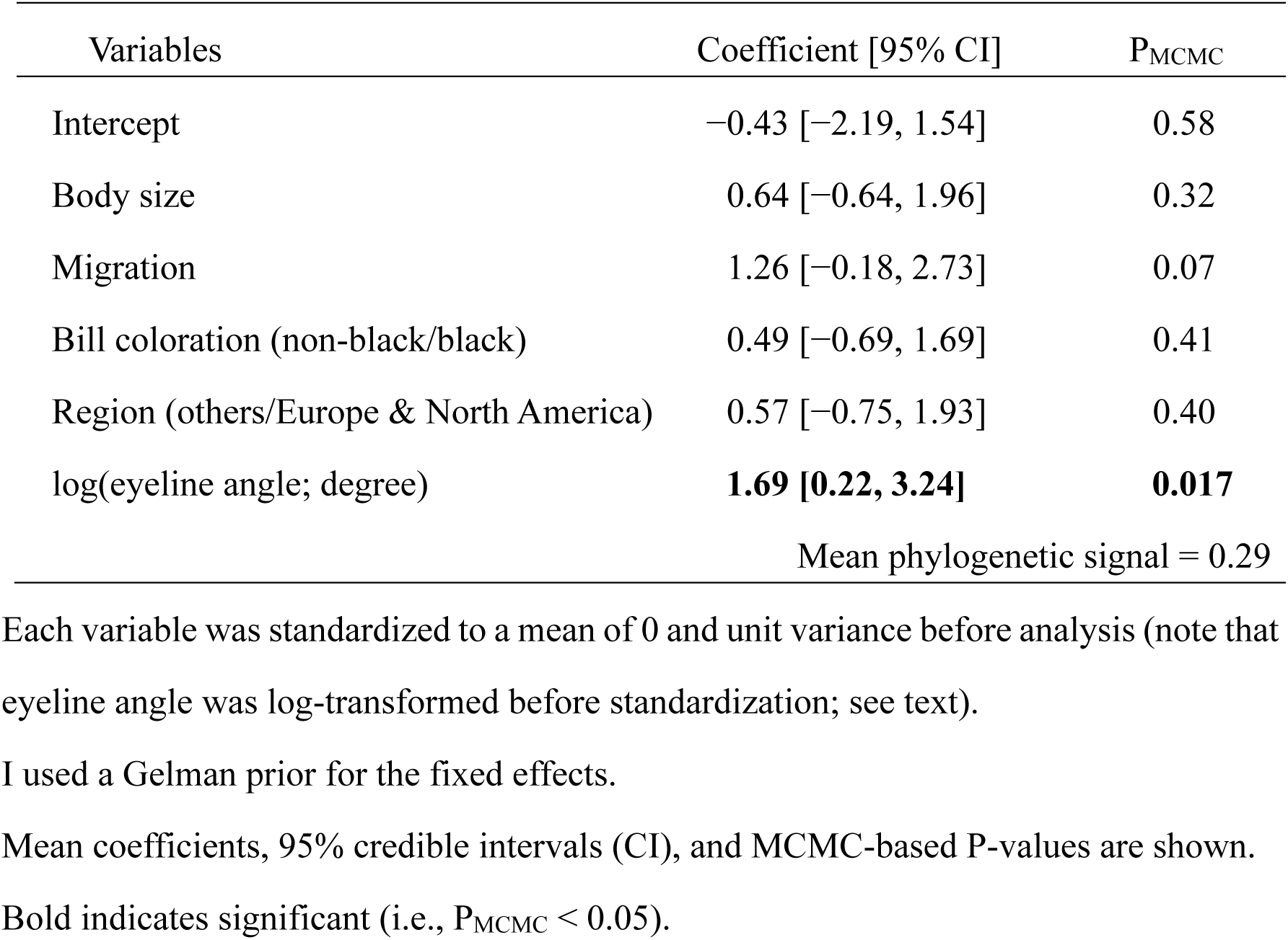
Multivariable Bayesian phylogenetic mixed model with a binomial error distribution predicting skimming in relation to eyeline angles in terns and allies (N = 40)

## Discussion

The current study demonstrated that tern lineages with foraging behaviors that require accurate aiming are more likely to have eyelines than others (Figs. 3 & 4), thereby supporting the sight-line hypothesis. This result corroborates the previous finding in hirundines, in which hirundine lineages foraging on large and active prey items (i.e., those need accurate aiming) are more likely to have eyelines than others (Hasegawa 2023). A positive association between body size and eyeline can also be explained by the sight-line hypothesis, because larger terns forage on larger (and thus swifter) prey than others (Gochfeld & Burger 1996), which requires more accurate aiming. Furthermore, among terns with eyelines, eyeline angle was evolutionarily associated with the occurrence of skimming behavior (i.e., a foraging behavior in which direction of travel is upwardly deviated from direction of bills and thus upwardly-tilted eyelines would be helpful; see Introduction). To my knowledge, this is the first study to demonstrate the key prediction of the sight-line hypothesis, i.e., direction of eyeline matters (Ficken et al. 1971), further supporting the hypothesis.

Previous studies of animal coloration often focus on color patches, and investigate its size and coloration in relation to survival, reproduction, and a variety of eco-evolutionary factors (e.g., see Hill & McGraw 2006 for a review on birds). This is not surprising, because, without selection on the size and coloration, the focal color patch would not evolve and be maintained. However, as shown here, “borderline,” i.e., color contrast between two adjacent colored regions (or, thin lines, in some cases), can have ecological functions and thus can evolve and be maintained under selection pressures (e.g., under selection for accurate aiming, here). In particular, direction of the borderlines is virtually ignored, although their importance was proposed more than fifty years ago (Fichen et al. 1971). The association between skimming and eyeline angle (Fig. 6) is difficult to explain with other explanations (e.g., upwardly-tilted eyelines would increase glare from the front side of the face and thus cannot be explained by simple glare reduction), thereby providing strong support for the sight-line hypothesis. Notably, the functional importance of the borderline does not preclude the functional importance of each color patch. Rather, selection on the borderline might promote the functional use of color patches (e.g., intra- and interspecific signaling, countershading), and vice versa. In fact, terns often have a black cap during the breeding period, which is absent during the nonbreeding period (e.g., the Whiskered tern, *Childonias hybrida*), indicating differential selection pressures between the two periods (e.g., sexual selection favors a black cap during the breeding season, whereas a black cap should be avoided in the nonbreeding period to avoid conflict with others: Bridge et al. 2005). As Fiche et al. (1971) already suggested, additional functions (e.g., sexual signaling) and the resultant color patterns would affect the function of sight-line and associated foraging behavior (skimming, here). Mutual functional (i.e., ecological) and evolutionary dependency of individual color patches and borderlines remains to be clarified.

A caveat of the current study is that, although I statistically controlled for phylogeny, body size, migratory habit, and bill coloration (Table 1; and potential research bias as well in Table 2), there might be unmeasured confounding factors. For example, the evolutionary association between the presence/absence of eyelines and foraging behavior might be explained by the third factor, such as countershading (e.g., Haney et al. 2020). However, contrasting dark–light patterns (rather than gradual change from dark to light coloration) does not fit the countershading pattern, and this explanation does not explain why dark–light borderlines occur just in front of the eyes (rather than lower or higher position). Moreover, qualitatively the same pattern was observed in two phylogenetically and ecologically different avian groups (i.e., terns and hirundines; see Introduction; also see Hasegawa 2023), and thus it is unlikely that confounding factors explain the two independent findings. Furthermore, an evolutionary association was observed between the direction of eyeline and foraging behavior (Table 2), which would be challenging to explain with the same confounding factors. Therefore, the current pattern, along with the previous one (Hasegawa 2023), fits the sight-line hypothesis rather than possible confounding factors, which should be tested by manipulative experiments in the future.

In conclusion, the current study provides macroevolutionary evidence that supports the sight-line hypothesis. The pattern of eyeline angle is particularly notable as it cannot be easily explained by other hypotheses (e.g., glare reduction; see above). The current study, however, is a correlational study and thus requires behavioral tests through manipulative experiments that can control unmeasurable confounding factors in the future. Although we often assume that animal coloration should have functions to catch (or, conversely, not to catch) the eyes of others (e.g., signaling, concealment), they might be used for catching their own eyes (i.e., aiming), which should not be ignored in future studies of animal coloration.

## Acknowledgments

I thank Dr Emi Arai for her comments and thank Dr Ichiro Tayasu and his lab members at Research Institute for Humanity and Nature for their valuable support. I also thank Dr Glaucia Del Rio and her lab members at Florida Museum of Natural History for their kindest support. This study was supported by the KAKENHI grant of the Japan Society for the Promotion of Science (JSPS: 25K09772).

## Author contribution

MH performed data analysis and wrote the manuscript.

## Ethical approval

This comparative study does not include any treatments of animals, as all the information was gathered from literatures.

## Data availability

The data sets supporting this article will be uploaded to osf.io.

## Supplementary Material 1

List of references used for searching for the occurrence of skimming behavior in terns and allies.

---

**Table S1.**
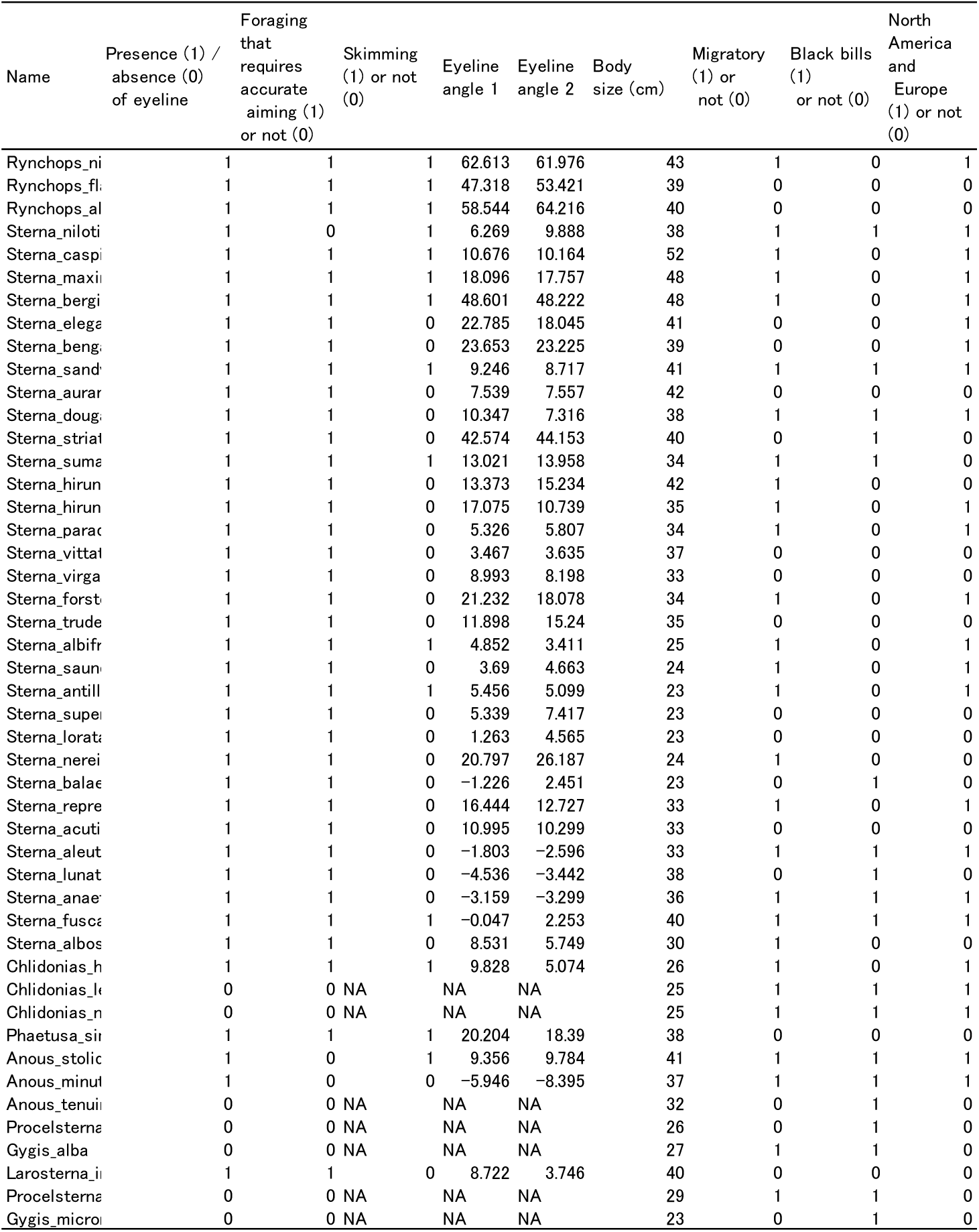
Dataset of the current study (will be attached just before submission).

**Fig. S1.**
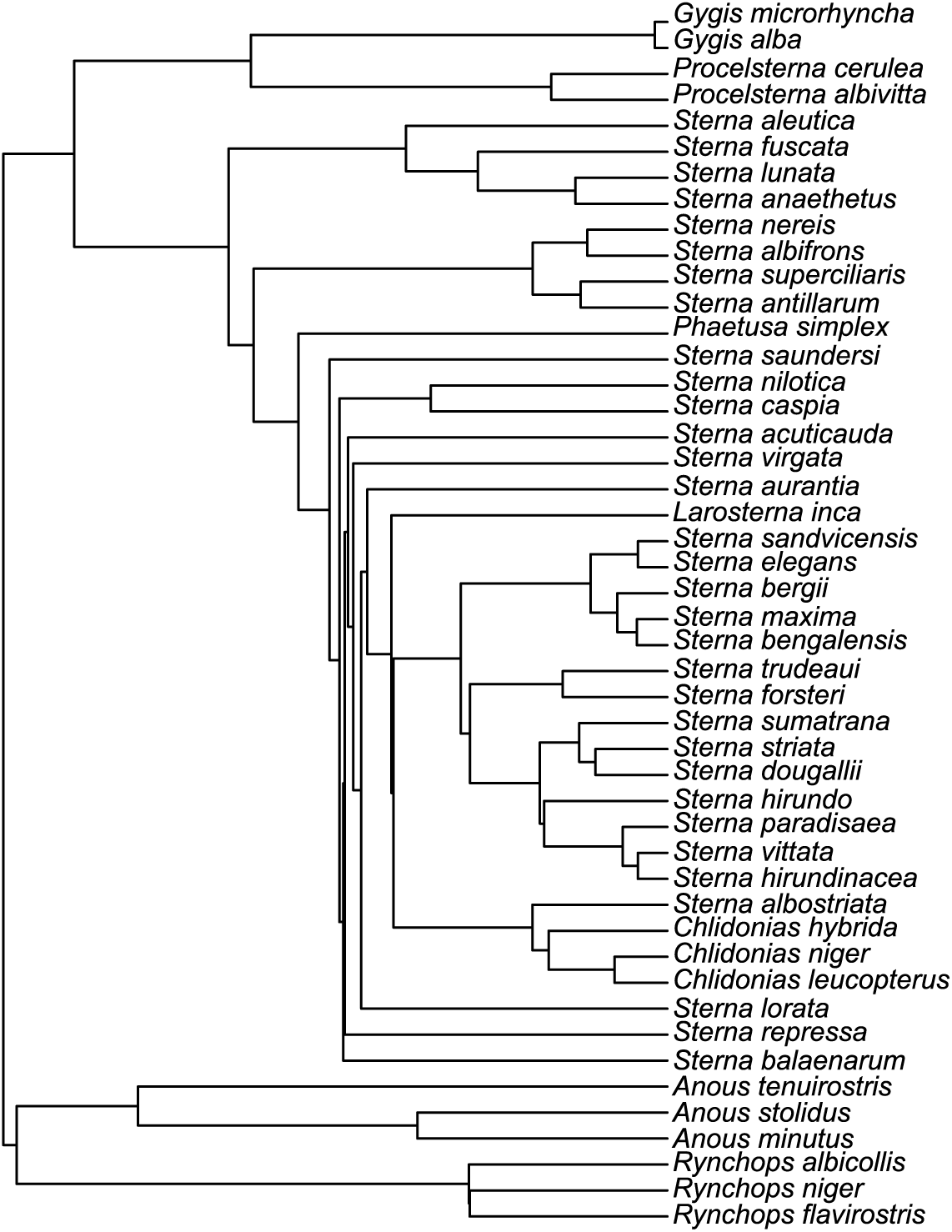
The least-squares consensus tree of terns using 1000 alternative trees obtained from birdtree.org (the function “ls.consensus” from the package “phytools”)

